# Deletion of CD226 in Foxp3^+^ T cells Reduces Diabetes Incidence in Non-Obese Diabetic Mice by Improving Regulatory T Cell Stability and Function

**DOI:** 10.1101/2022.06.02.494443

**Authors:** Puchong Thirawatananond, Matthew E. Brown, Lindsey K. Sachs, Juan M. Arnoletti, Wen-I Yeh, Amanda L. Posgai, Melanie R. Shapiro, Yi-Guang Chen, Todd M. Brusko

## Abstract

Co-stimulation serves as a critical checkpoint for T cell development and activation, and several genetic variants affecting co-stimulatory pathways confer risk for autoimmune diseases. A single nucleotide polymorphism in *CD226* (*rs763361*; G307S) has been shown to increase susceptibility to type 1 diabetes, multiple sclerosis, and rheumatoid arthritis. CD226 competes with the co-inhibitory receptor TIGIT (T cell immunoreceptor with Ig and ITIM domains) to bind CD155 to amplify TCR signaling. We previously found that *Cd226* knockout protected non-obese diabetic (NOD) mice from disease, but the impact of CD226 signaling on individual immune subsets remained unclear. We focused on regulatory T cells (Tregs) as a population of interest, as prior reports demonstrated that human CD226^+^ Tregs exhibit reduced FOXP3^+^Helios^+^ purity and suppressive function following expansion. Hence, we hypothesized that global deletion of *Cd226* would increase Treg stability and accordingly, Treg-specific *Cd226* deletion would inhibit diabetes in NOD mice. Indeed, crossing the NOD.*Cd226*^*-/-*^ and NOD.*Foxp3*-GFP-Cre.*R26*-loxP-STOP-loxP-YFP Treg-fate tracking strains resulted in increased Treg induction and decreased FoxP3-deficient “ex-Tregs” in the pancreatic lymph nodes. We generated a Treg-conditional knockout (Treg^Δ*Cd226*^) strain and found that female Treg^Δ*Cd226*^ mice had decreased insulitis and diabetes incidence compared to Treg^WT^ mice. Additionally, we observed increased TIGIT expression on Tregs and conventional CD4^+^ T cells within the pancreas of Treg^Δ*Cd226*^ versus Treg^WT^ mice. These findings demonstrate that an imbalance of CD226/TIGIT signaling may contribute to Treg destabilization in the NOD mouse and highlight the potential for therapeutic targeting of this pathway to prevent or reverse autoimmunity.

## Introduction

Co-stimulatory signaling is essential for the development and activation of T lymphocytes, and gene variants converging on several co-stimulatory pathways confer risk for autoimmune diseases, including type 1 diabetes (T1D). The *rs763361* (C>T) single nucleotide polymorphism (SNP) within the *CD226* gene, for example, has been shown to increase susceptibility to T1D, multiple sclerosis (MS), and rheumatoid arthritis (RA) (1, 2). CD226, also known as DNAX accessory molecule-1 (DNAM-1), is expressed on natural killer (NK) cells (3), CD8^+^ T cells (4), platelets (5), monocytes (6, 7), and activated CD4^+^ T cells (8). CD226 competes with the co-inhibitory receptors TIGIT (T cell immunoreceptor with Ig and ITIM domains) and CD96 to bind the ligand CD155 (poliovirus receptor, PVR) on antigen presenting cells (APCs) (9). Upon ligation, CD226 initiates phosphorylation of its cytoplasmic tail leading to downstream PI3K/Akt and MAPK/Erk signaling, and inducing T cell activation and proliferation (3). The autoimmune-associated Ser^307^ variant, encoded by the *rs763361* (T) allele, is thought to potentially provide an additional cytoplasmic phosphorylation site, augmenting downstream signaling (10). Hence, there is a need to understand how CD226 activity may contribute to the pathogenesis of T1D and other autoimmune diseases.

CD4^+^ regulatory T cells (Treg) play an important role in T1D, with the majority of T1D immunotherapies seeking to augment Treg function or frequency relative to effector T cell subsets (11, 12). We previously reported that IFN-γ producing Tregs were enriched in CD226 expression (13). However, how CD226 signaling affects Treg stability and function during T1D development remains unclear. We previously reported that expanded human CD226^+^ Tregs had increased expression of interferon-γ as well as decreased suppression of CD4^+^ and CD8^+^ T cell responders compared to CD226^-^ Tregs (14). Building on this, we observed increased FOXP3^+^Helios^+^ purity and *in vitro* suppressive capacity in CD4^+^CD25^+^CD226^-^ versus CD4^+^CD25^+^CD127^lo/-^ sorted Tregs, supporting the potential utility of CD226 as a negative selection marker for improved sorting prior to Treg adoptive cell therapy (15). Consistent with these observations, Sato et al. recently identified CD226 signaling as a potential source of Treg instability, finding that pre-treatment of human peripheral blood mononuclear cells (PBMCs) with an α-human CD226 blocking antibody diminished graft versus host disease (GvHD) development in a humanized mouse model by increasing human CD4^+^FOXP3^+^ Treg numbers and upregulating their FOXP3 expression (16).

We previously investigated the causal role of CD226 signaling in autoimmune diabetes through the genomic knockout (gKO) of *Cd226* in the non-obese diabetic (NOD) mouse model of T1D (17). Genetic deletion of *Cd226* led to reduced diabetes incidence and insulitis severity. Additionally, this germline deletion was associated with increased CD8^+^ T cell output from the thymus but decreased CD44^+^CD62L^-^ memory CD8^+^ T cells in the pancreatic lymph nodes (PLN) and islet-specific glucose-6-phosphatase catalytic subunit related protein (IGRP)-specific memory CD8^+^ T cells in the pancreas. Despite our findings suggesting that *Cd226* gKO primarily impacted the pathogenicity of CD8^+^ T cells, adoptive transfer of mixed populations of *Cd226*^*+/+*^ and *Cd226*^*-/-*^T cells into immunodeficient NOD.*scid* recipients demonstrated a requirement for CD226 on CD4^+^ T cells to confer disease (17). In line with our findings, Wang and colleagues demonstrated that CD226 gKO delayed disease onset in the experimental autoimmune encephalomyelitis (EAE) mouse model of MS (18). Interestingly, *Cd226*^*-/-*^ EAE mice also had increased amounts of Tregs and increased induced Treg (iTreg) proliferation *in vitro* (18), and α-CD226 blocking antibodies reduced Th1-mediated autoimmunity (8) and increased IL-10 production (19) in the EAE mouse. While these observations collectively support the role of CD226 in promoting the loss of T cell tolerance in NOD and EAE mice, it was unclear whether the influence of CD226 on Treg proliferation and stability would extend to the NOD mouse model.

The current study investigates how CD226 signaling impacts Treg frequency and function in the context of the NOD mouse model. We hypothesized that Treg-selective *Cd226* KO would improve Treg stability, resulting in reduced diabetes incidence in these mice. To test this, we generated two novel NOD strains, specifically a Treg-fate reporter (20) *Cd226* gKO and a Treg *Cd226* conditional KO (cKO) strain, for detailed interrogation of Treg-specific contributions toward insulitis and autoimmune diabetes.

## Materials and Methods

### Animal Husbandry and Regulatory Approval

Animals were bred and housed in specific-pathogen free facilities at the University of Florida with *ad libitum* access to food and water. Studies were performed with protocols approved by the University of Florida Institutional Animal Care and Use Committee (UF IACUC) and in accordance with the National Institutes of Health Guide for Care and Use of Animals.

### Mouse Strains

NOD.*Foxp3*-GFP-Cre.*R26*-loxP-STOP-loxP-YFP Treg-fate tracking mice (20) were crossed with our previously reported NOD.*Cd226*^-/-^ (17) strain to generate Treg-fate reporter mice with *Cd226* gKO on the NOD background. The *Cd226* cKO allele was generated at Jackson Laboratories through CRISPR/Cas9-mediated knock-in of loxP within intronic regions flanking exon 2 of the *Cd226* gene on the NOD background. The following single guide RNAs were used: intron 1, GCTATCGTACAGGGATGGAT and GCTTTGCTATCGTACAGGGA; intron 2, TGACATGTGCAGGATCTCTT and GAAATACTAAAATTTACAAG. Founder cKO mice with confirmed loxP knock-ins were backcrossed with NOD mice for one generation before crossing with NOD.*Foxp3*-GFP-Cre mice purchased from Jackson laboratories (20). *Cd226*^fl/+^.*Foxp3*^Cre/+^ x *Cd226*^fl/+^.*Foxp3*^+/+^ breeder pairs were created and, subsequently, *Cd226* Treg-specific cKO (i.e., *Cd226*^*fl/fl*^ x *Cd226*^*fl/fl*^) and *Cd226* intact (*Cd226*^*+/+*^ x *Cd226*^*+/+*^) breeding schemes were utilized to obtain Treg^Δ*Cd226*^ and Treg^WT^ mice for experiments.

### Genotyping

The *Cd226* gKO allele genotyping was performed with a custom Taqman SNP assay (Thermo Fisher Scientific) using the following primers: forward, GGGAGCATGAAGAGCATCCT; reverse, GCGACATCTGTAAAGTCCTGAGT; *Cd226* WT reporter (VIC), CAAATGCCATGGCCGCT; *Cd226* gKO reporter (FAM), CAAATGCCATGCCGCT (17). The *Foxp3*-GFP-Cre transgene was genotyped using a SYBR Green PCR Master Mix (Thermo Fisher Scientific) to amplify the GFP allele (forward, GACCCTGAAGTTCATCTGCACC; reverse, CGGGTCTTGTAGTTGCCGTC) and internal control (IC; forward, GGCAAAGGTGGAAATGAAGA; reverse, CTCAGACCACACAGGGAATG). A Roche LightCycler 480 was used to detect GFP and IC amplicons at melting points of 86ºC and 79ºC, respectively. The *Cd226 fl* allele was genotyped using high resolution melting by amplifying with SYBR Green PCR Master Mix (Thermo Fisher Scientific; forward, CTGGCACAGAGGACACACTC; reverse, GCACAGGAAAGAAGTTTCAGC) on a Roche LightCycler 480. Samples were assessed based on melting profiles and difference plots for *Cd226 fl* (approximately 200 base pairs) versus *Cd226* WT alleles (approximately 180 base pairs).

### DNA confirmation of Treg *Cd226* cKO

Splenocytes from mice were stained with α-CD4 (PerCP-Cy5.5; clone GK1.5; BioLegend) and α-CD8α (BV711; 53-6.7; BioLegend) and CD8^+^ T cells, CD4^+^GFP^-^ Tconv and CD4^+^GFP^+^ Treg were sorted on a FACSAria III (BD Biosciences). Sorted cells were then processed with a DNeasy kit (Qiagen), and 2 ng of DNA was amplified with the following primers complementary to *Cd226* gene sequences flanking exon 2: forward: TTGTTGCACAGAGCTAAGTCTG, reverse: CCACCTGGGTTAAGTTATGCG. Samples were then run on a 1.5% agarose (w/v) gel with a 100-base pair ladder (Thermo Fisher Scientific).

### Flow Cytometry

12-week-old euglycemic *Cd226* WT, gKO, Treg^Δ*Cd226*^, and Treg^WT^ mice were humanely euthanized, and thymus, spleen, MLN, PLN, and pancreas were harvested. Thymus and spleens were homogenized using a syringe plunger over a 40 μm filter while PLNs were homogenized using two frosted glass microscope slides. Pancreas tissues were minced into 1 mm pieces and incubated in complete RPMI [cRPMI; 50 μg/mL penicillin/streptomycin, 2 mM Glutamax (Life Technologies), 20 mM sodium pyruvate (Gibco), 5 mM non-essential amino acids (Gibco), 5 mM HEPES (Gibco), 10% fetal bovine serum (Biofluid), 20 mM NaOH (Sigma-Aldrich), 50 mM 2-mercaptoethanol (Sigma Aldrich)] with 1 mg/mL collagenase IV (Gibco) and 1 U/mL DNase I (Thermo Scientific) for 18 minutes at 37°C. Following digestion, pancreas single-cell suspensions were washed with cRPMI and strained through a 40 μm filter. Cells were stained with Live/Dead Near-Infrared dye (L/D NIR; Invitrogen) according to the manufacturer’s instructions, then washed with stain buffer. Cells were resuspended in Brilliant Violet Stain Buffer Plus (BD Biosciences) with anti-CD16/32 Fc-Block (BioLegend), then stained with the following antibodies for 30 minutes at room temperature: α-CD3ε (BV605; clone 17A2; BioLegend), α-CD4 (PerCP-Cy5.5; GK1.5; BioLegend), α-CD8α (BV711; 53-6.7; BioLegend), α-CD44 (PE; IM7; BioLegend), α-CD62L (APC; MEL-14; BioLegend), α-CD226 (BV650; TX42.1; BioLegend), α-TIGIT (PE-Dazzle594; 1G9; BioLegend), and α-CD183 (PE-Cy7; CXCR3-173; BioLegend). Cells were washed with stain buffer, then fixed and permeabilized with the eBioscience FOXP3 Transcription Factor Staining kit (Invitrogen) according to the manufacturer’s instructions. Cells were then stained with α-T-bet (BV421; 4B10; BioLegend) overnight prior to acquisition on a Cytek Aurora 5-laser spectral flow cytometer. The flow cytometric panel assessing thymic Treg frequency included α-CD4 (PerCP-Cy5.5; GK1.5; BioLegend), α-CD8α (BV711; 53-6.7; BioLegend), α-CD44 (PE; IM7; BioLegend), α-CD226 (BV650; TX42.1; BioLegend), and α-Nrp1 (BV421; 3E12; BioLegend). The intracellular flow cytometry panel to quantify Treg frequency included α-CD4 (PerCP-Cy5.5; GK1.5; BioLegend), α-CD8α (BV711; 53-6.7; BioLegend), α-CD44 (PE; IM7; BioLegend), α-CD226 (BV650; TX42.1; BioLegend), and α-Helios (Pacific Blue; 22F6; BioLegend). Data were analyzed using FlowJo v10 software (TreeStar).

### Insulitis and Dacryoadenitis Scoring

Pancreata from Treg^Δ*Cd226*^ and Treg^WT^ mice were fixed overnight in 10% formalin in PBS, then washed twice with PBS and stored in 70% ethanol until further processing. Fixed pancreata were paraffin embedded, sectioned at 250 μm steps, stained with hematoxylin and eosin (H&E), and imaged with an Aperio Scancope CS slide scanner. Three sections of pancreas per mouse were scored (mean of islets per mouse: 59.7, standard deviation: 12.4) for insulitis severity by a blinded observer according to previously published criteria (21). Briefly, a score of 0 indicates no infiltration, 1 indicates peri-insulitis, 2 indicates <50% islet immune infiltration, and 3 indicates >50% islet immune infiltration (21). Lacrimal gland inflammatory scoring was performed by quantifying foci (i.e., ≥50 mononuclear cells) and normalizing to 4mm^2^ glandular surface area as previously described (22).

### Diabetes Incidence Study

Blood glucose measurements from Treg^Δ*Cd226*^, Treg^WT^ and NOD.*Cd226*^*fl/fl*^.*Foxp3*^*+/+*^ mice were taken weekly beginning at 7 weeks of age by collecting a small drop of blood (<5μL) onto an AlphaTRAK 2 glucometer via tail vein prick. Animals registering blood glucose >250 mg/dL had a confirmatory screening 24 hours later, and two consecutive readings >250 mg/dL indicated the onset of diabetes. Mice were monitored until diabetes onset or up to 30 weeks of age, at which time they were humanely euthanized.

### Phosflow

Splenocytes from Treg^Δ*Cd226*^ and Treg^WT^ mice were plated in a 96-well plate at a concentration of 1,000,000 cells/mL with plate-bound α-CD3ε (2 μg/mL) and plate-bound CD155 (1 μg/mL; BioLegend) for 15 or 30 minutes at 37ºC. Stimulations were halted with BD Cytofix Fixation Buffer (BD Biosciences) followed by a 10-minute incubation at 37ºC and wash with PBS. Cells were then stained with L/D NIR for 15 minutes and quenched with stain buffer. Fixed cells were permeabilized with 90% methanol for 1 hour and washed with stain buffer twice. Permeabilized cells were incubated with anti-CD16/32 Fc-Block (BioLegend) and stained for CD4 (PerCP-Cy5.5; GK1.5; BioLegend), CD8α (BV711; 53-6.7; BioLegend), p-S6 (PE; D57.2.2E; Cell Signaling Technology), and pAkt (AF647; 193H12; Cell Signaling Technology) overnight at 4ºC. Samples were then acquired on a Cytek Aurora 5-laser spectral flow cytometer, and data were analyzed using FlowJo v10 software (TreeStar).

### *In Vitro* Proliferation Assay

Splenocytes from Treg^Δ*Cd226*^ and Treg^WT^ mice were stained with CellTrace Violet proliferation dye (CTV; Invitrogen) and quenched with cold cRPMI as per manufacturer instructions. Cells were plated in a 24-well plate at a concentration of 500,000 cells/mL on either plate-bound α-CD3ε (2 μg/mL; 145-2C11; BioLegend) and soluble α-CD28 (1 μg/mL; 37.51; BioLegend) or plate-bound α-CD3ε (2 μg/mL) and plate-bound CD155-Fc (1 μg/mL; BioLegend) for four days. Cells were then harvested, incubated with anti-CD16/32 Fc-Block (BioLegend), and stained with L/D NIR, α-CD4 (PerCP-Cy5.5; GK1.5; BioLegend), and α-CD8α (BV711; 53-6.7; BioLegend) as described above and acquired on a Cytek Aurora 5-laser spectral flow cytometer. Data were analyzed using FlowJo v10 software (TreeStar).

### Statistical Analyses

Statistical analysis was performed with GraphPad Prism version 9 (GraphPad Software). Flow cytometry data were analyzed using multiple t-tests with Sidak-Bonferroni multiple comparisons correction, 1-way ANOVA with Bonferroni’s multiple comparisons test, and 2-way ANOVA with Sidak multiple comparisons correction. Insulitis scoring was compared by Chi-square test. Diabetes incidence curves were compared by Log-rank (Mantel-Cox) test. *P* values < 0.05 were considered significant.

## Results

### CD226 is upregulated on ex-Treg and effector/memory Treg subsets

Previous studies have shown that expression of CD226 on murine conventional CD4^+^ T cells (Tconv) is upregulated during initial T cell receptor (TCR) activation and maintained following Th1 skewing (8). We hypothesized that CD226 would be similarly upregulated in activated effector Tregs. Indeed, our analysis of previously published human RNA-seq datasets (23) revealed that *CD226* transcription was increased in activated Tregs compared to resting Tregs (24). To confirm that this observation extended to protein expression levels in mice, we performed flow cytometry to assess CD226 levels on splenocytes from the Treg-fate tracking NOD strain, NOD.*Foxp3*-GFP-Cre.*R26*-loxP-STOP-loxP-YFP. In this strain, Tregs express a GFP-Cre recombinase product driven by a *Foxp3* promoter-driven transgene, which facilitates the permanent expression of YFP under the constitutively active *Rosa26* promoter after excision of a floxed stop codon preceding the YFP transgene. This would provide the ability to discern GFP^+^YFP^-^ “early” Tregs which are thought to be Tregs that recently induced Foxp3 expression, GFP^+^YFP^+^ stable Tregs, and GFP^-^YFP^+^ ex-Tregs that have lost Foxp3 expression. The investigators that developed this strain found that ex-Tregs were enriched for a CD44^+^CD62L^-^ effector-memory (EM) phenotype and led to hyperglycemia and pancreatic islet destruction during adoptive transfers into NOD *Rag2*^*-/-*^ recipients (20); hence, we predicted that ex-Tregs would have upregulated CD226 levels compared to “early” Tregs and stable Tregs. As expected, CD226 surface expression was increased in the ex-Treg compartment compared to that of early Tregs (1.4-fold; p=0.0289), CD44^-^CD62L^+^ naïve Tregs (2.6-fold; p<0.0001), and CD44^+^CD62L^-^ EM Tregs (1.4-fold; p<0.0001) (**Figure 1A-B & Supplemental Figure 1A**). In contrast, surface expression of TIGIT was increased in ex-Tregs compared to early Tregs (1.5-fold; p=0.0194) and naïve Tregs (1.4-fold; p<0.0001), but was comparable to EM Tregs (p=0.1576) (**Figure 1C-D**). In addition to the studies demonstrating the role of CD226 in disrupting FOXP3 expression (16) while TIGIT restores Treg suppression (25), this data suggests that an imbalance of CD226:TIGIT signaling might play a role in ex-Treg development, contributing to the loss of Foxp3 expression in EM Tregs and promoting the proliferation of ex-Tregs.

**Figure 1.**
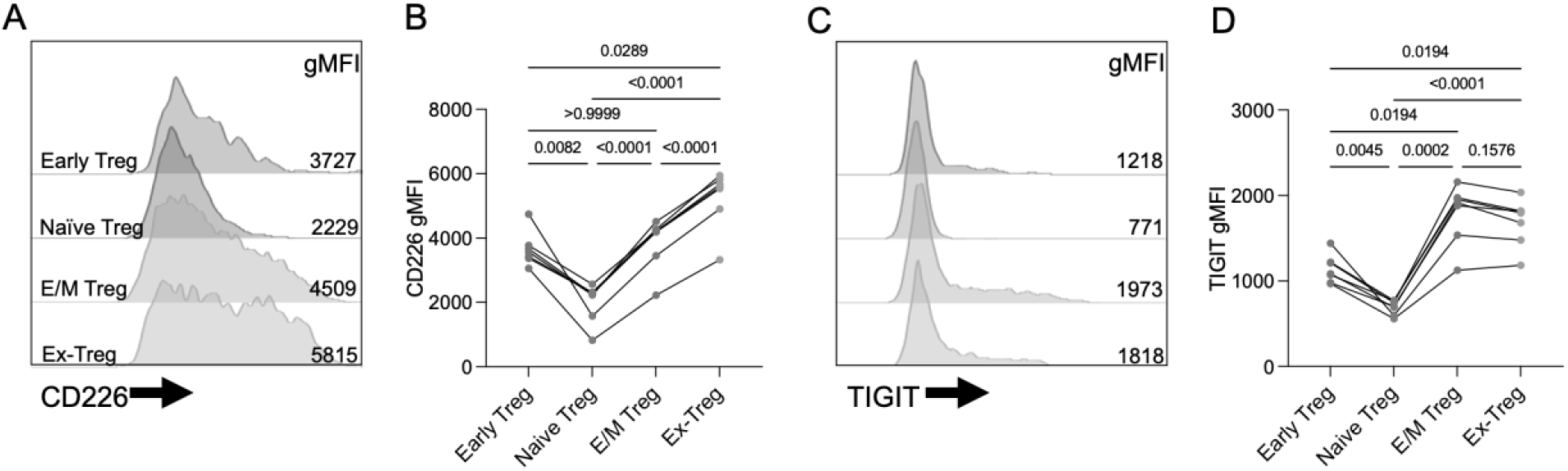
CD226 and TIGIT expression levels in Tregs and ex-Tregs. Splenocytes from pre-diabetic 12-week-old male Treg-fate tracking NOD mice were assessed through spectral flow cytometry. **(A)** Half-offset histograms and **(B)** quantification of CD226 geometric mean fluorescence intensity (gMFI) on GFP^+^YFP^-^ early Treg, GFP^+^YFP^+^CD44^-^CD62L^+^ naïve Treg, GFP^+^YFP^+^CD44^+^CD62L^-^ EM Treg, and GFP^-^YFP^+^ ex-Treg subsets. **(C)** Half-offset histograms and **(D)** quantification of TIGIT gMFI. *P* values are displayed and were determined from repeated measures one-way ANOVA with Bonferroni’s multiple comparisons test (*n*=7).

### *Cd226* gKO reduced ex-Treg frequency in pancreatic lymph nodes of NOD mice

To determine the impact of *Cd226* KO on Treg stability in the NOD mouse, we crossed our NOD.*Cd226* gKO strain (17) with Treg-fate tracking mice and assessed Treg frequencies in various tissues from 12-week-old euglycemic *Cd226*^*+/+*^ wild-type (WT) and *Cd226*^*-/-*^ gKO offspring through flow cytometry. Interestingly, frequencies of early (GFP^+^YFP^-^), normal (GFP^+^YFP^+^), and ex-Tregs (GFP^-^YFP^+^) were not significantly altered between *Cd226* WT and *gKO* mice in the thymus, spleen, and mesenteric lymph nodes (**Figure 2A-C**). This suggests that germline loss of CD226 signaling did not impair thymic Treg output to peripheral organs. However, the PLNs of *Cd226* gKO mice had increased early Treg frequencies (9.7-fold; *p*=0.0002) but decreased normal Treg (0.37-fold; *p*=0.0016) and ex-Tregs (0.45-fold; *p*=0.0011) compared to those of *Cd226* WT mice (**Figure 2A-D**). This suggests that CD226 signaling might restrain Foxp3 induction while promoting Foxp3 instability in the context of an inflammatory microenvironment such as the PLN in NOD mice. Based on the opposing roles of CD226 and TIGIT in immune cell activation, we assessed the level of TIGIT expression and found that while there was no difference in TIGIT expression in the thymus, spleen, or MLN, there was increased expression on Tconv (1.55-fold; *p*=0.0317) and CD8^+^ T cells (1.59-fold; *p*=0.0450), but unchanged on Tregs (including early, normal, and ex-Treg subsets), in PLNs from *Cd226* gKO versus WT mice (**Supplemental Figure 1B-H**). This suggests that loss of CD226 coupled with increased TIGIT may contribute to the diabetes protection previously observed in *Cd226* gKO NOD mice (17).

**Figure 2.**
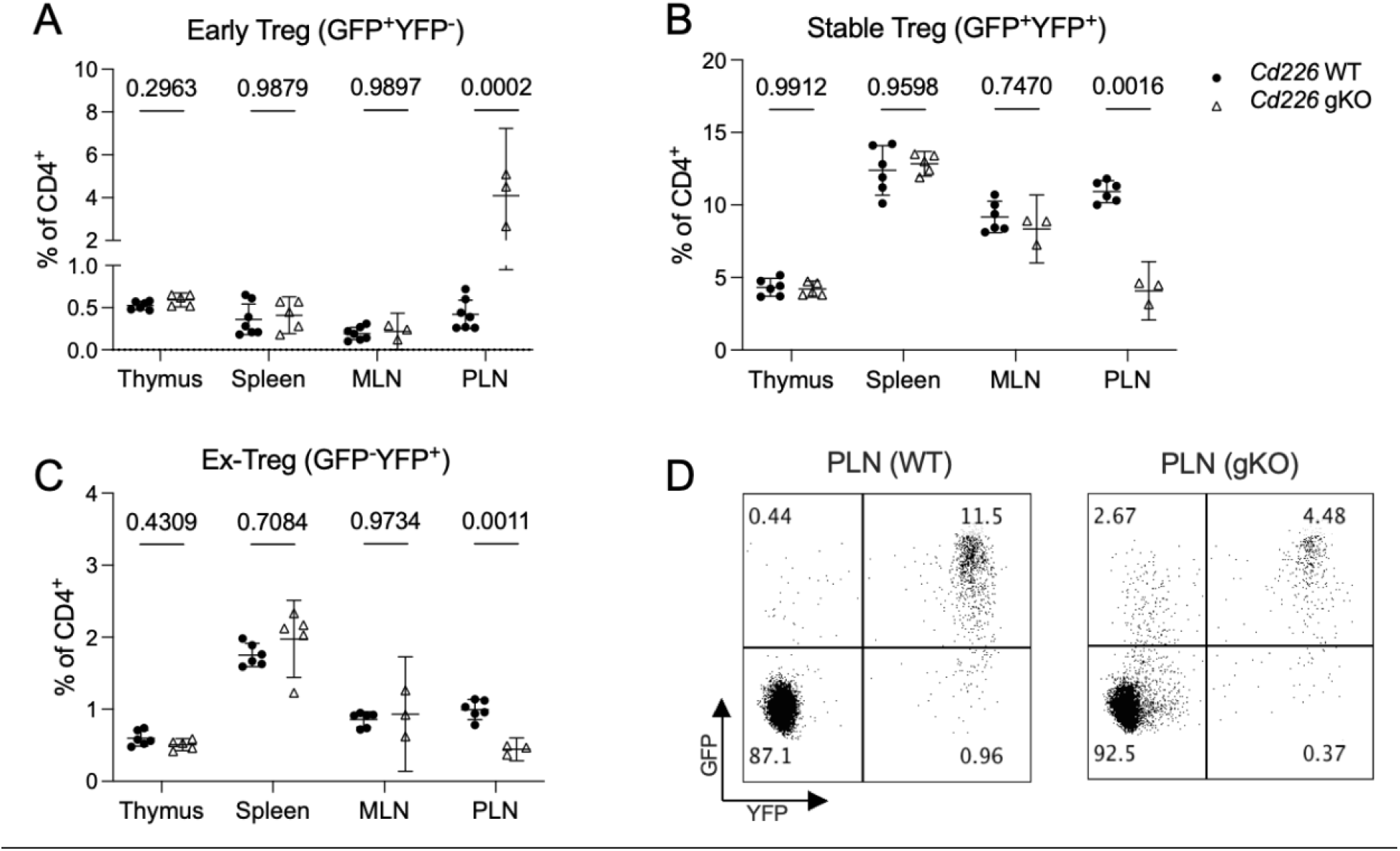
Decreased stable and ex-Tregs and increased pTreg in PLN of CD226 gKO NOD mice. **(A)** GFP^+^YFP^-^ “early” Tregs, **(B)** GFP^+^YFP^+^ stable Tregs, and **(C)** GFP^-^YFP^+^ ex-Tregs were quantified from thymus, spleen, mesenteric LN (MLN) and PLN from 12-week-old pre-diabetic male Treg-fate reporter mice with *Cd226* WT (circles, *n*=7) or global knockout (triangles, gKO; *n*=5) alleles (plotted as a % of CD4^+^ T cells following singlet, live, CD3ε^+^ gating; analyzed using multiple t-tests with Sidak-Bonferroni multiple comparisons correction). **(D)** Representative flow cytometry from PLN of *Cd226* WT and gKO mice.

### Generation of *Cd226*^*fl/fl*^ Treg-specific cKO NOD mice

While the germline *Cd226* KO demonstrated a reduction of ex-Treg, an increase in early Tregs, and an enrichment for TIGIT^+^ Tconv and CD8^+^ T cells in the disease-relevant inflammatory environment of the PLN, it does not preclude the possibility that these observations could be secondary to the diabetes-protected phenotype of the *Cd226* gKO, resulting from Treg-extrinsic factors (17). To investigate the intrinsic role of CD226 on Tregs and how the loss of this signaling might affect diabetes pathogenesis, we generated NOD mice expressing a *Cd226* cKO allele by inserting loxP sequences flanking exon 2 of *Cd226* (*Cd226*^*fl/fl*^). We then crossed these mice with the NOD.*Foxp3*-GFP-Cre strain (20) to selectively reduce CD226 expression in Foxp3-expressing Tregs (**Fig. 3A**). PCR amplification of the *Cd226*^*fl*^ allele in sorted CD4^+^GFP^+^ Treg, CD4^+^GFP^-^ Tconv, and CD8^+^ T cells from *Cd226*^*fl/fl*^.*Foxp3*-GFP-Cre^+/-^ (Treg^Δ*Cd226*^) splenocytes showed a truncated *Cd226* amplicon (381 base pairs) from Tregs compared to CD4^+^ Tconv and CD8^+^ T cells (868 base pairs) (**Supplemental Figure 2A**), confirming successful cKO. Expression of CD226 was reduced on Tregs in a *Cd226*^*fl*^ allele-dose dependent manner (**Fig. 3B-C**). Inhibition of CD226 expression was strongly selective towards Tregs, as CD226 surface expression was maintained in CD4^+^ Tconv and CD8^+^ T cells (**Fig. 3D-E**). Overall, these data validate the selectiveness of the Treg^Δ*Cd226*^ strain, leading us to study the impacts of this genetic modification on diabetes pathogenesis.

**Figure 3.**
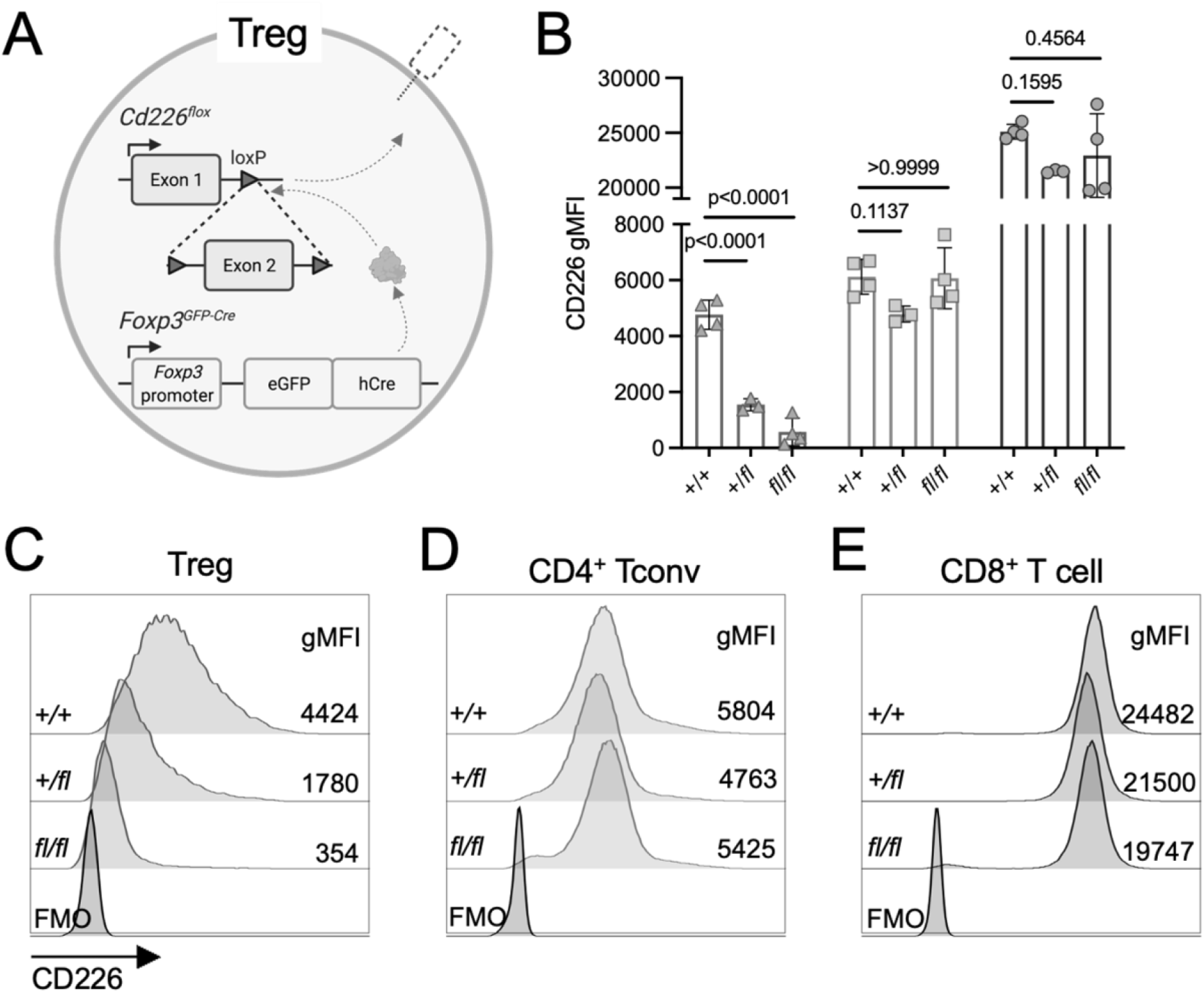
CD226 gMFI is selectively reduced in Treg cKO NOD mice. **(A)** Schematic of loxP insertions flanking exon 2 (*fl*) of *Cd226* and excision in Tregs of Treg^Δ*Cd226*^ mice. **(B)** Scatter plot showing CD226 gMFI and **(C-E)** representative histograms of CD4^+^GFP^+^ Tregs (triangles), CD4^+^GFP^-^ Tconv (squares), and CD8^+^ T cells (circles) of 12-week-old pre-diabetic female *Cd226* +/+ (*n*=4), +/fl (*n*=3), and fl/fl (*n*=3) mice versus fluorescence minus one (FMO) control. Analyzed with one-way ANOVA with Bonferroni’s multiple comparisons test.

### Treg *Cd226* cKO reduced insulitis and diabetes incidence in female NOD mice

To determine the effect of the selective deletion of CD226 on Tregs in the context of diabetes pathogenesis in NOD mice, we performed insulitis scoring on hematoxylin and eosin (H&E)-stained pancreas sections from age-matched pre-diabetic Treg^Δ*Cd226*^ and Treg^WT^ mice (**Fig. 4A-B**). Insulitis scores were reduced in 12-week-old female Treg^Δ*Cd226*^ mice (*p*<0.0001) while insulitis severity was comparable in 16-week-old male Treg^Δ*Cd226*^ and Treg^WT^ NODs (*p*=0.47, **Fig. 4C-D**). In line with this, female Treg^Δ*Cd226*^ mice had reduced diabetes incidence (44.0%) as compared to Treg^WT^ (66.7%; *p*=0.042) and *Cd226*^*fl/fl*^.*Foxp3*^*+/+*^ controls (75.0%; *p*=0.012). We did not observe a significant difference in diabetes incidence between female Treg^WT^ and *Cd226*^*fl/fl*^.*Foxp3*^*+/+*^ control groups (*p*=0.5985). Interestingly, diabetes incidence in male mice was unchanged between Treg^Δ*Cd226*^ and Treg^WT^ genotypes (*p*=0.3367, **Fig. 4E-F**). While we previously reported that *Cd226* gKO led to decreased dacryoadenitis in male NOD mice (17), we did not observe a significant difference in lacrimal gland focus scores of 16-week-old Treg^WT^ and Treg^Δ*Cd226*^ mice (*p*=0.1613; **Supplemental Figure 2B-D**). Thus, we chose to focus on female NOD mice in our further characterization of the Treg^Δ*Cd226*^ strain.

**Figure 4.**
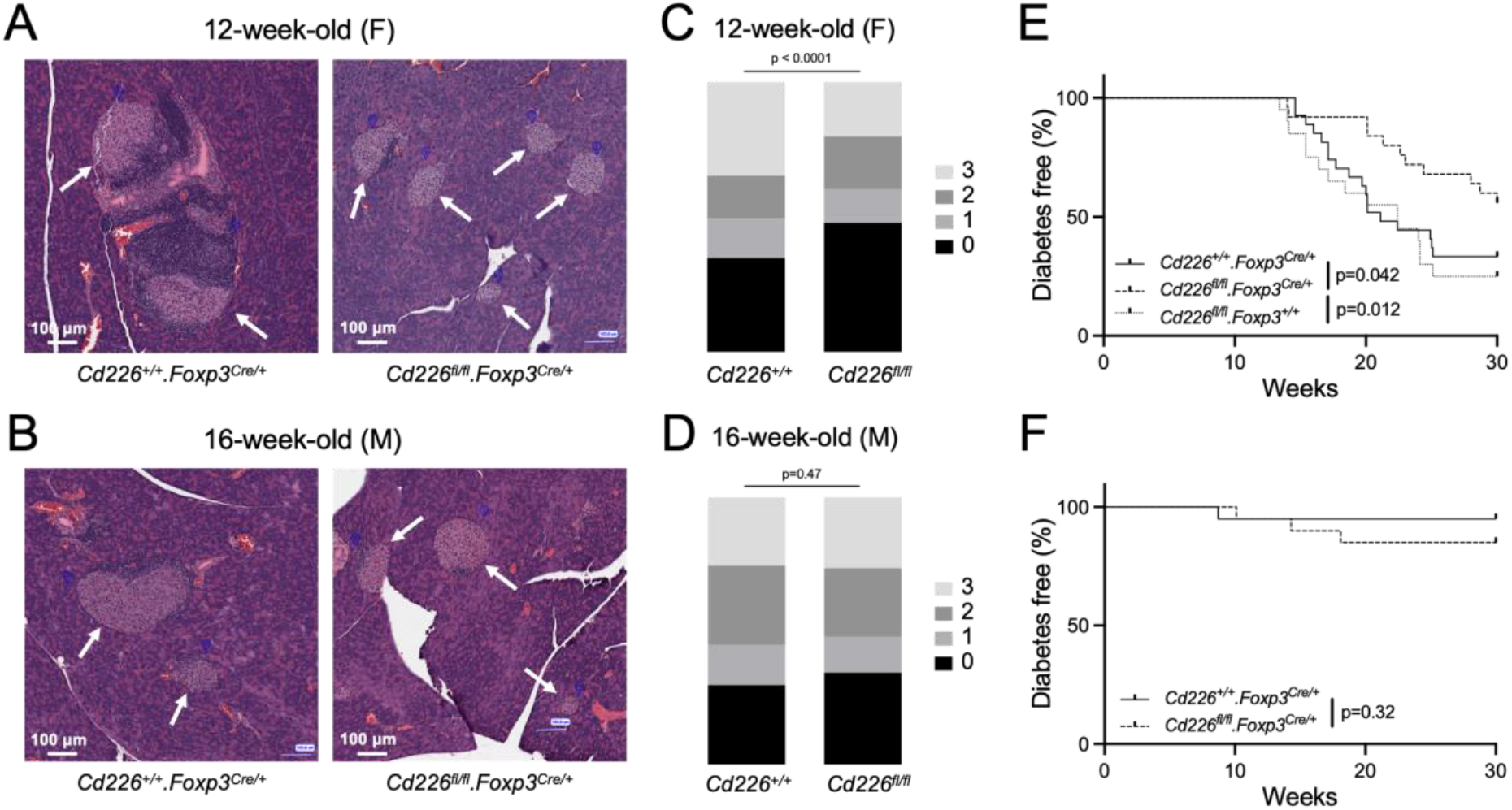
Treg *Cd226* cKO reduces insulitis and diabetes incidence in female NOD mice. **(A-B)** Representative H&E-stained pancreas sections with white arrows denoting islets of Langerhans and **(C-D)** stacked bar graphs depicting insulitis scores from 12-week-old female and 16-week-old male *Cd226*^+/+^.*Foxp3*^*Cre/+*^ (female, *n*=8 mice and 512 islets; male, *n*=5 mice and 252 islets) and *Cd226*^*fl/fl*^ .*Foxp3*^*Cre/+*^ (female, *n*=8 mice and 443 islets; male, *n*=3 mice and 262 islets) NOD mice. Statistical analysis performed with Chi-square test. **(E-F)** Diabetes incidence curves from **(E)** female and **(F)** male NOD mice. Female *N* = *Cd226*^+/+^.*Foxp3*^*Cre/+*^, 27; *Cd226*^*fl/fl*^.*Foxp3*^*Cre/+*^, 20; *Cd226*^*fl/fl*^.*Foxp3*^*+/+*^, 20. Male *N* = *Cd226*^+/+^.*Foxp3*^*Cre/+*^, 20; *Cd226*^*fl/fl*^.*Foxp3*^*Cre/+*^, 20. Analyzed with Log-rank (Mantel-Cox) test.

### TIGIT surface expression is increased on Tregs in Treg^Δ*Cd226*^ mice

To determine the basis of the protection observed in female Treg^Δ*Cd226*^ mice, we first performed *ex vivo* phenotyping of spleen, PLN, and pancreas by flow cytometry (**Supplemental Figure 3A**). We observed that bulk CD8^+^ and CD4^+^ T cell frequencies were unchanged in the spleen, PLN, and pancreas of Treg^Δ*Cd226*^ and Treg^WT^ mice (**Supplemental Figure 3B-C**). Within the CD4^+^ compartment, CD4^+^Foxp3^-^ Tconv and CD4^+^Foxp3^+^ Treg frequencies were similar between Treg^Δ*Cd226*^ and Treg^WT^ NODs in all three tissues evaluated (**Supplemental Figure 3D-E**). Since CD226 is upregulated in EM T cell subsets from WT NODs (**Figure 1**) (8, 14), we next examined the frequencies of CD44^-^CD62L^+^ naïve and CD44^+^CD62L^-^ EM T cells and found that there was no difference in naïve or EM CD8^+^ T cells, CD4^+^ Tconv and Tregs between Treg^Δ*Cd226*^ and Treg^WT^ mice (**Supplemental Figure 3F-K**). Furthermore, Treg^Δ*Cd226*^ and Treg^WT^ mice showed no differences in thymic Treg (Foxp3^+^Helios^+^ or Foxp3^+^Nrp1^+^) or peripheral Tregs (Foxp3^+^Helios^-^ or Foxp3^+^Nrp1^-^) frequencies in the thymus, spleen, PLN, and pancreas (**Supplemental Figure 4**) (26-29). Overall, this data suggests that the protection from diabetes provided by Treg^Δ*Cd226*^ was not due to increased numbers of Tregs in the periphery and that Treg *Cd226* cKO did not lead to any intrinsic changes in memory Treg formation or extrinsic changes in Tconv cells.

Due to the opposing roles of CD226 and TIGIT and the increased TIGIT on effector T cells observed in *Cd226* gKO PLNs (**Supplemental Figure 1H**), we next examined TIGIT geometric mean fluorescence intensity (gMFI) in Treg^Δ*Cd226*^ mice. Interestingly, TIGIT surface expression on Tregs was increased in the spleens (*p*=0.0002), PLNs (*p*=0.0002), and pancreas (*p*<0.0001) of Treg^Δ*Cd226*^ mice (**Fig. 5A-B**). In contrast, there was increased TIGIT expression in CD4^+^ Tconv in the spleen (*p*=0.0461) and pancreas (*p*<0.0001), but not PLN (*p*=0.2501), of Treg^Δ*Cd226*^ mice (**Fig. 5C**). Furthermore, there was no change in TIGIT surface expression in CD8^+^ T cells in the organs examined (**Fig. 5D**). This suggests that increased TIGIT signaling in Tregs secondary to the loss of CD226 may contribute to the delayed disease progression observed in female Treg^Δ*Cd226*^ NOD mice.

**Figure 5.**
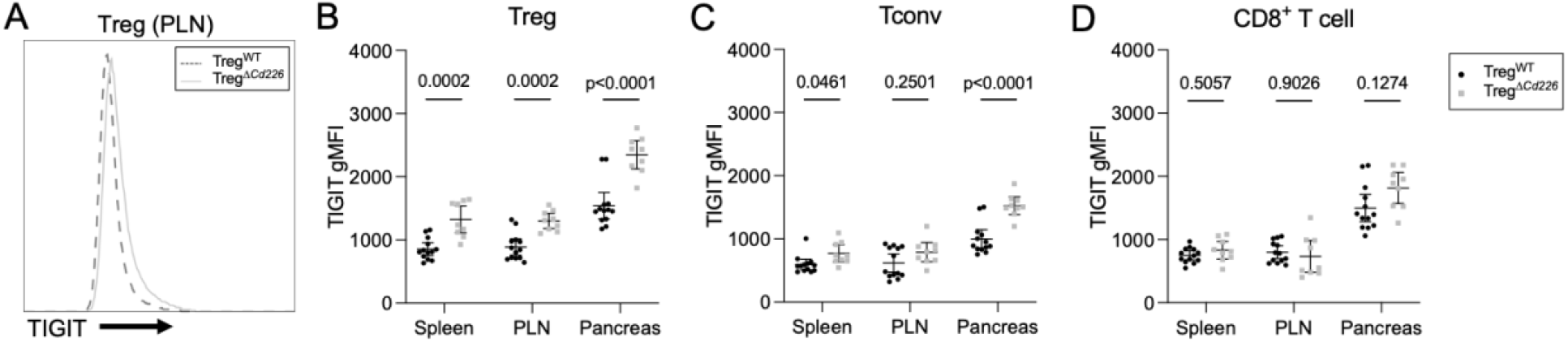
Treg^Δ*Cd226*^ mice show increased TIGIT expression compared to Treg^WT^ mice. **(A)** Overlaid histograms of TIGIT surface expression on Tregs in the PLN showing Treg^WT^ (dashed) and Treg^Δ*Cd226*^ (solid). Quantification of TIGIT gMFI of **(B)** Tregs, **(C)** CD4^+^ Tconv, and **(D)** CD8^+^ T cells from 12-week-old female Treg^WT^ (circles) and Treg^Δ*Cd226*^ (squares) mice. Statistical analysis was performed with multiple t-tests with Sidak-Bonferroni multiple comparisons correction.

### Treg^Δ*Cd226*^ effector T cells show reduced proliferation in response to *in vitro* stimulation

To assess whether the lack of CD226 signaling on Tregs would affect T cell activation, we performed short time-course stimulation studies to determine the magnitude of TCR and co-stimulatory signaling via measurement of phosphorylated Akt (pAkt; pS473) and ribosomal protein S6 (pS6; pS235/pS236) in splenocytes activated with plate-bound α-CD3ε/CD155-Fc. We found that while there was no difference in pAkt gMFI of Treg^Δ*Cd226*^ and Treg^WT^ splenocytes (**Fig. 6A-C**), there was a decrease in pS6^+^ CD8^+^ T cells from Treg^Δ*Cd226*^ mice after 30 minutes of stimulation (*p*=0.0304; **Fig. 6D-F**). This suggests that Treg *Cd226* cKO may result in a CD8^+^ T cell-extrinsic mechanism of TCR signaling. We next stimulated splenocytes *in vitro* with α-CD3ε/α-CD28 or α-CD3ε/CD155-Fc conditions and assessed the proliferation of T cell subsets via dye dilution as measured by flow cytometry. In response to α-CD3ε/CD155-Fc, but not α-CD3ε/α-CD28, proliferation was reduced in CD8^+^ (*p*=0.0030) and CD4^+^ T cells (*p*=0.0002) from Treg^Δ*Cd226*^ compared to Treg^WT^ mice (**Fig. 6G-J**). Altogether, these findings suggest that reduced CD8^+^ T cell proliferation, secondary to reduced S6 phosphorylation downstream of TCR signaling, could be a key mechanism by which insulitis and diabetes incidence are inhibited in Treg^Δ*Cd226*^ mice.

**Figure 6.**
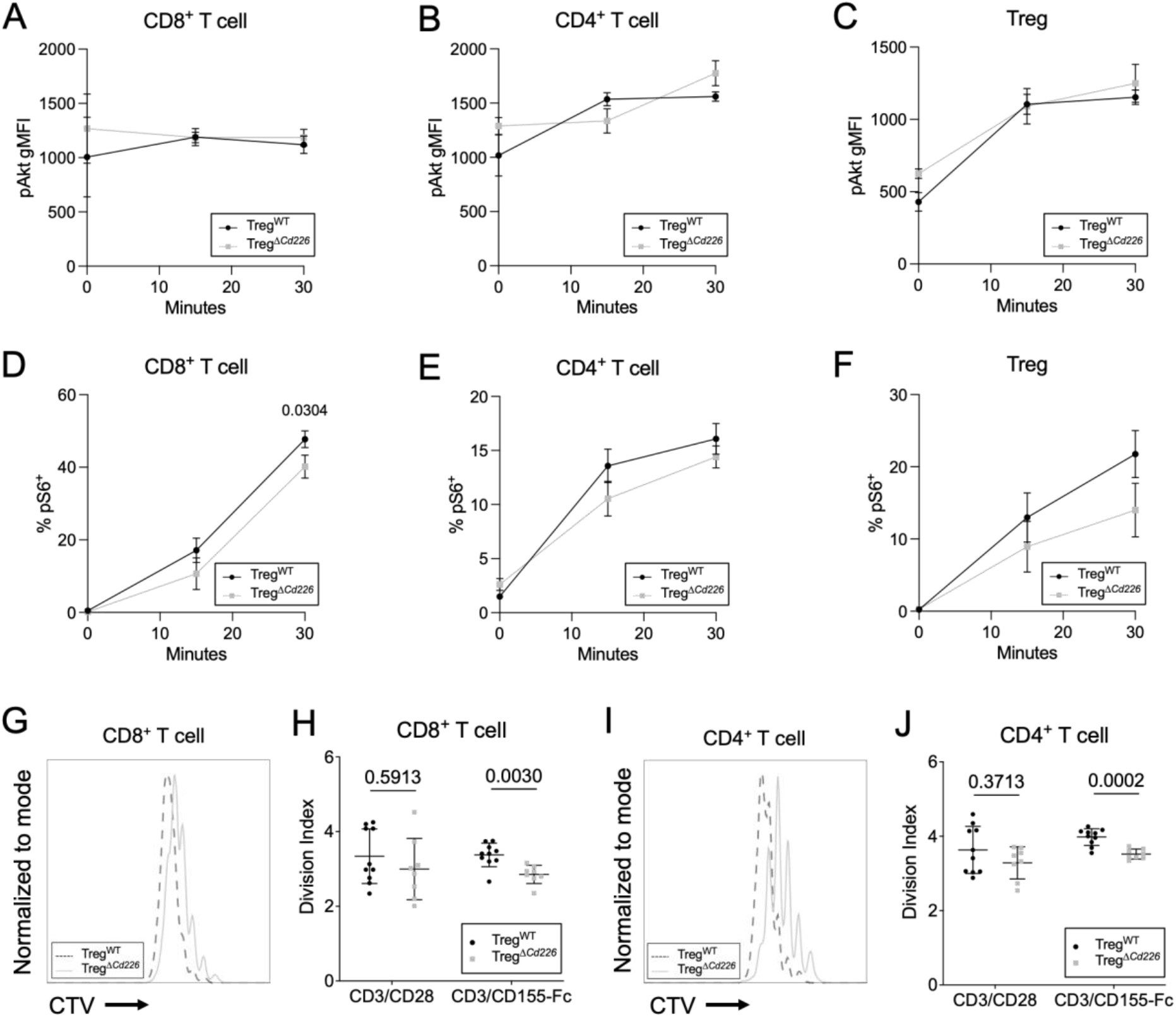
Treg^Δ*Cd226*^ splenocytes have reduced proliferation in response to α-CD3ε/CD155-Fc stimulus. Quantification of **(A-C)** phospho-Akt (pAkt) gMFI and **(D-F)** phospho-S6^+^ (pS6) ribonuclear protein in CD8^+^ T cells, CD4^+^ T cells, and Treg from Treg^WT^ (circles) and Treg^Δ*Cd226*^ (squares) splenocytes (*n*=4) after stimulation with plate-bound α-CD3ε/CD155-Fc. Statistical analyses were performed with 2-way ANOVA with Sidak multiple comparisons correction. **(G-J)** Overlaid histograms showing CellTrace Violet dye dilution and division indices of CD8^+^ **(G** and **H)** and CD4^+^ T cells **(I** and **J)** from 12-week-old female Treg^WT^ (dashed/black; *n*=10) and Treg^Δ*Cd226*^ (solid/square; *n*=7) splenocytes after four days of stimulation with plate-bound α-CD3ε/α-CD28 or plate-bound α-CD3ε/CD155-Fc. Analyzed with multiple t-tests with Sidak- Bonferroni multiple testing correction.

## Discussion

In this study, we determined that CD226 signaling in Tregs contributes to diabetes pathogenesis in female NOD mice. In agreement with previous reports (23, 24), we found that stable EM Tregs highly expressed CD226. Additionally, we found significantly higher CD226 expression in ex-Tregs which has not been previously reported. Global *Cd226* deletion on the NOD background led to decreased ex-Treg frequencies in the PLN, highlighting the notion that excess activation provided by CD226 signaling may lead to Treg instability and the loss of Foxp3 expression. In contrast, others have shown in a humanized mouse model of GVHD that murine Tregs deficient in CD226 have greater TIGIT signaling and in turn, improved maintenance of Foxp3 expression (16). Interestingly, *Cd226* gKO did not affect early Treg, normal Treg, or ex-Treg frequencies in the thymus, spleen, or MLN, demonstrating that CD226 deletion did not impair central or peripheral Treg development. This is in contrast with the defects observed in NOD.*Cd28* KO mice in thymic Treg development and peripheral maintenance when the loss of CD28-B7 co-stimulation resulted in decreased CD25 expression on Tregs and IL-2 production by Tconv (30-32). The protection and reduced PLN ex-Tregs in *Cd226* gKO mice prompted us to interrogate the role of CD226 on Treg function in NOD disease apart from the risk conferred through CD226 activity within other immune cell subsets.

Generation of the NOD.*Cd226*^*fl/fl*^ strain and crossing with the NOD.*Foxp3*-GFP-Cre strain demonstrated Treg-selective *Cd226* cKO and decreased diabetes pathogenesis in female NOD mice. Flow cytometric studies showed that naïve and EM Treg, CD4^+^ Tconv, and CD8^+^ T cell frequencies were unaffected in Treg^Δ*Cd226*^ mice, suggesting there was no impairment of T cell activation of Treg cKO mice. We also observed a significant increase in TIGIT surface expression on Tregs in the spleen, PLN, and pancreas, along with increased TIGIT expression of CD4^+^ Tconv in the pancreas of Treg^Δ*Cd226*^ versus Treg^WT^ mice. Wang and colleagues observed a similar phenotype, reporting that *Cd226*^-/-^ C57BL/6 mice after inducing EAE showed increased CTLA-4 and TIGIT expression on Tregs in the central nervous system (33). We observed decreased pS6^+^ in CD8^+^ T cells and proliferation of CD4^+^ and CD8^+^ T cells in response to *in vitro* α-CD3ε/CD155-Fc stimulation of Treg^Δ*Cd226*^ versus Treg^WT^ splenocytes. We surmise that the inhibition of Teff proliferation in Treg^Δ*Cd226*^ mice may be due to enhanced Treg suppression mediated through increased TIGIT-CD155 interactions, which others have shown to increase IL-10 production by CD4^+^ T cells and Tregs (34, 35) as well as dendritic cells (36). Decreased S6 phosphorylation, which lies downstream of the Akt signaling cascade, corresponds with a previous report showing that the *Cd226*^*-/-*^ EAE mouse had decreased pAkt and pErk signaling in Tregs (33). This suggests that CD226-deficient Tregs prevent diabetes pathogenesis by reducing effector T cell activation and proliferation through increased IL-10 production mediated by increased TIGIT-CD155 interactions.

The delayed onset and reduced incidence of diabetes in genomic and Treg-conditional *Cd226* KO NOD mice demonstrates the utility of the NOD model in studying interventions modulating CD226/TIGIT signaling and how they may affect autoimmune T1D pathogenesis. The interesting finding that there was no difference in insulitis or diabetes progression in male Treg^Δ*Cd226*^ mice could be due to the inherently low disease incidence of WT male NOD mice (5% in the current study) (37). Future studies may include crossing Treg-fate and *Cd226* cKO strains to determine how CD226 activity in Tregs intrinsically affects their potential development into ex-Tregs. Additionally, the NOD.*Cd226*^*fl/fl*^ mice could readily be crossed with additional strains, such as NOD.*Gzmb*-Cre and NOD.*Cd8*-Cre mice, to study effects of NK and CD8^+^ T cell-specific *Cd226* cKO. The detailed mechanistic studies reported herein support strategies to block CD226 signaling, particularly on Tregs, to interrupt T1D progression. Hence, future studies are needed to evaluate therapeutics modulating this costimulatory pathway for the ability to delay or prevent autoimmune diabetes in the NOD pre-clinical model in support of future translation to clinical trials.

## Acknowledgments

We would like to acknowledge Thinzar Myint, M.S. (University of Florida) for project coordination and support. We would also like to thank Kayla Nguyen and Jin-Ju Lee (University of Florida) for technical assistance with blood glucose monitoring. Treg-fate tracking NOD mice were a kind gift from Dr. Jeffrey Bluestone (University of California San Francisco).

## Author Contributions

PT: Data curation, Formal analysis, Funding acquisition, Investigation, Methodology, Project administration, Validation, Visualization, Writing-original draft, Writing-review & editing

MEB: Investigation, Visualization, Writing-review & editing LKS: Investigation, Writing-review & editing

JMA: Investigation, Writing-review & editing W-IY: Investigation, Writing-review & editing ALP: Writing-review & editing

MRS: Funding acquisition, Investigation, Methodology, Writing-review & editing YGC: Investigation, Writing-review & editing

TMB: Conceptualization, Funding acquisition, Methodology, Resources, Supervision, Writing-review & editing

