## Supplementary Figures for "Deletion of CD226 in Foxp3^+^ T cells Reduces Diabetes Incidence in Non-Obese Diabetic Mice by Improving Regulatory T Cell Stability and Function"

### Supplemental Figures

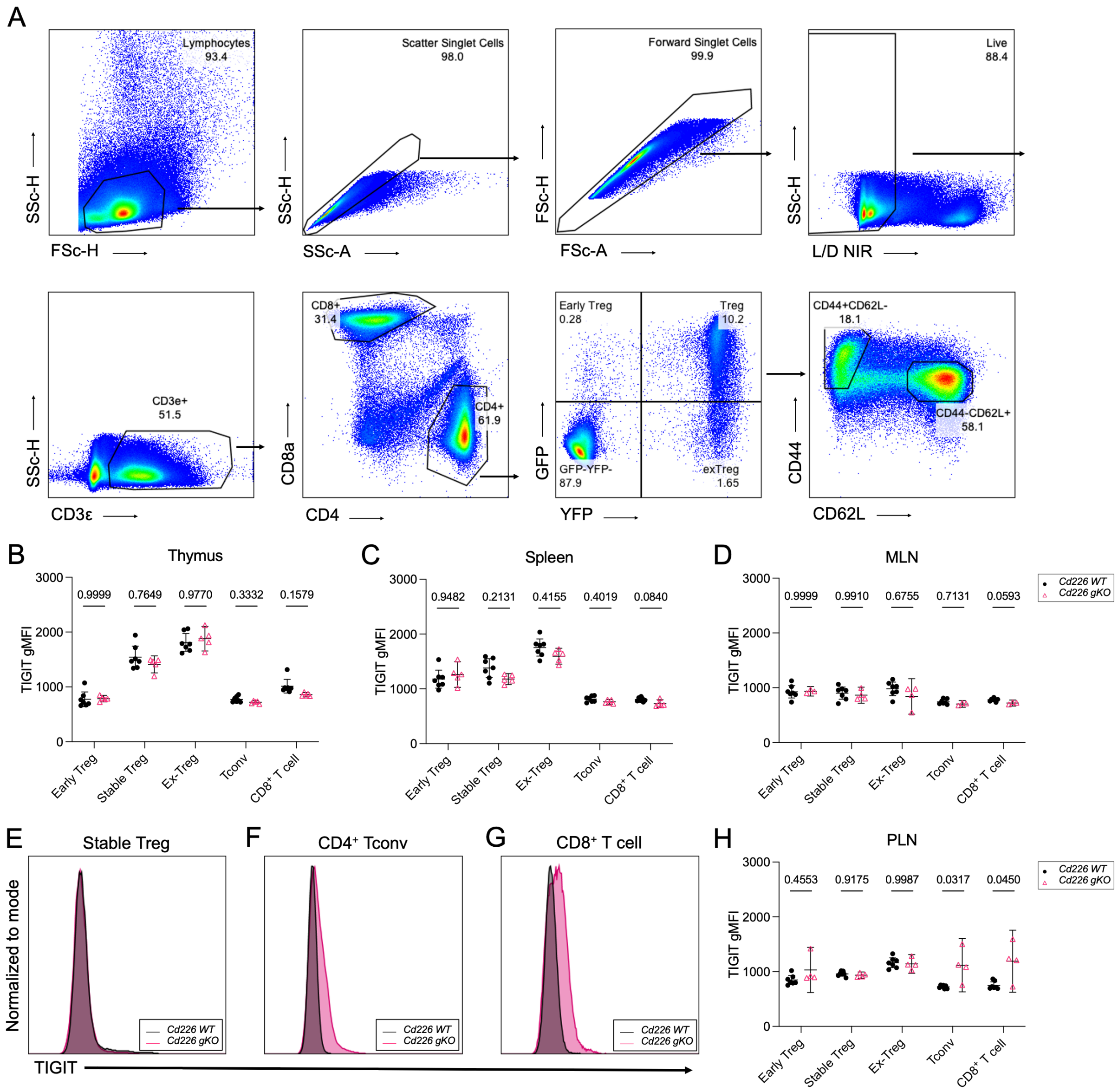


**Supplemental Figure 1. TIGIT levels are increased in CD4$^+$ Tconv and CD8$^+$ T cells in the PLNs of *Cd226* gKO NOD mice. (A)** Sample gating scheme for Treg-fate reporter mouse tissues analyzed on a 5-laser Cytek Aurora. Percentages of parent gates are listed underneath the gate labels. Representative data are shown from spleen. Quantification of TIGIT gMFI on GFP$^+$YFP$^-$ early Treg, GFP$^+$YFP$^+$ stable Treg, GFP$^-$YFP$^+$ ex-Treg, CD4$^+$ Tconv, and CD8$^+$ T cell subsets in the **(B)** thymus, **(C)** spleen, and **(D)** MLNs of 12-week-old male *Cd226* WT (black) and *Cd226*

gKO (pink) mice. Overlaid histograms of TIGIT gMFI of **(E)** stable Tregs, **(F)** $CD4^+$ Tconv, and **(G)** $CD8^+$ T cells in the PLNs of *Cd226* WT (black) and *Cd226* gKO (pink) mice with the quantification shown in **(H)**. Statistical analysis was performed with multiple t-tests with Sidak-Bonferroni multiple testing correction. (WT, *n*=7; gKO, *n*=5).

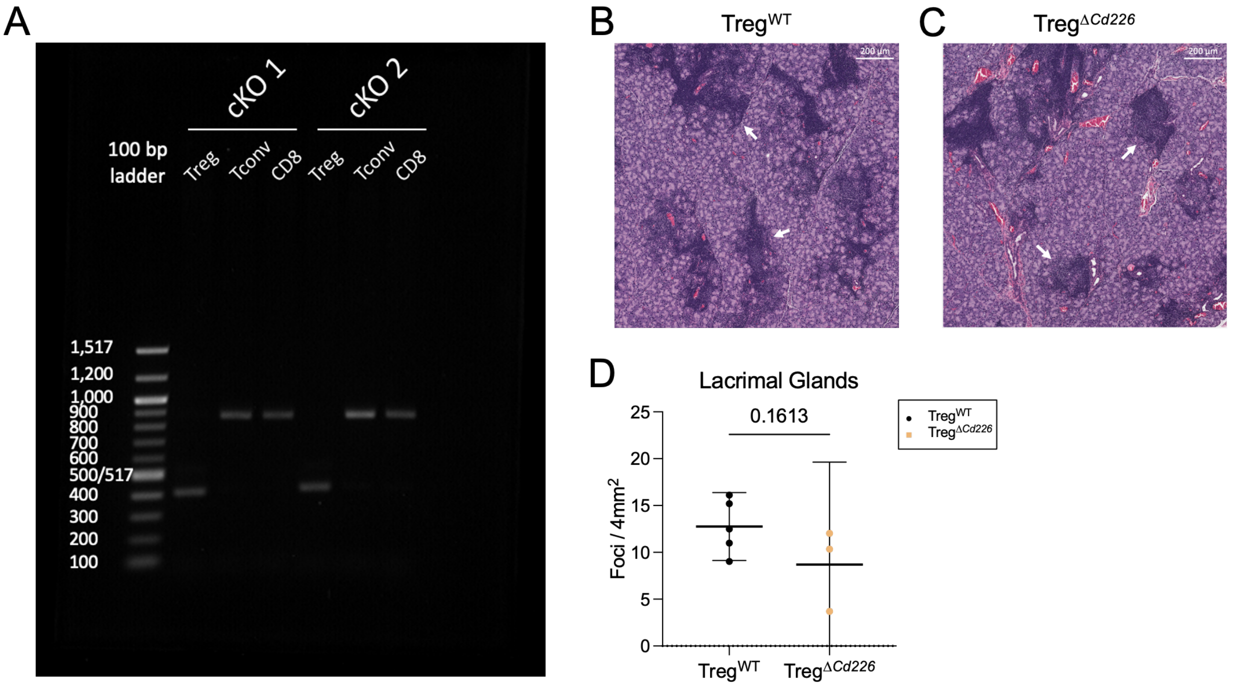


**Supplemental Figure 2. *Cd226* cKO results in gene truncation and does not affect NOD dacryoadenitis.** (A) $CD4^+GFP^+$ Tregs, $CD4^+GFP^-$ Tconv, and $CD8^+$ T cells were sorted from splenocytes from two conditional KO (cKO) mice on a BD FACSAria III. Genomic DNA was isolated from sorted T cell populations, and PCR was performed using primers amplifying the region flanking the flox insertions surrounding exon 2 of the *Cd226* gene. PCR amplicons were run on a 1.5% agarose gel with a 100-base pair (bp) DNA ladder. **(B-C)** Representative H&E-stained sections of lacrimal glands from 16-week-old male **(B)** Treg$^{WT}$ and **(C)** Treg$^{\Delta Cd226}$ mice with inflammatory foci labeled with white arrows. **(D)** Quantification of foci scoring (number foci per 4 mm$^2$) in the lacrimal glands. Unpaired t-tests were performed for statistical analysis (Treg$^{WT}$, $n$=4; Treg$^{\Delta Cd226}$, $n$=3).

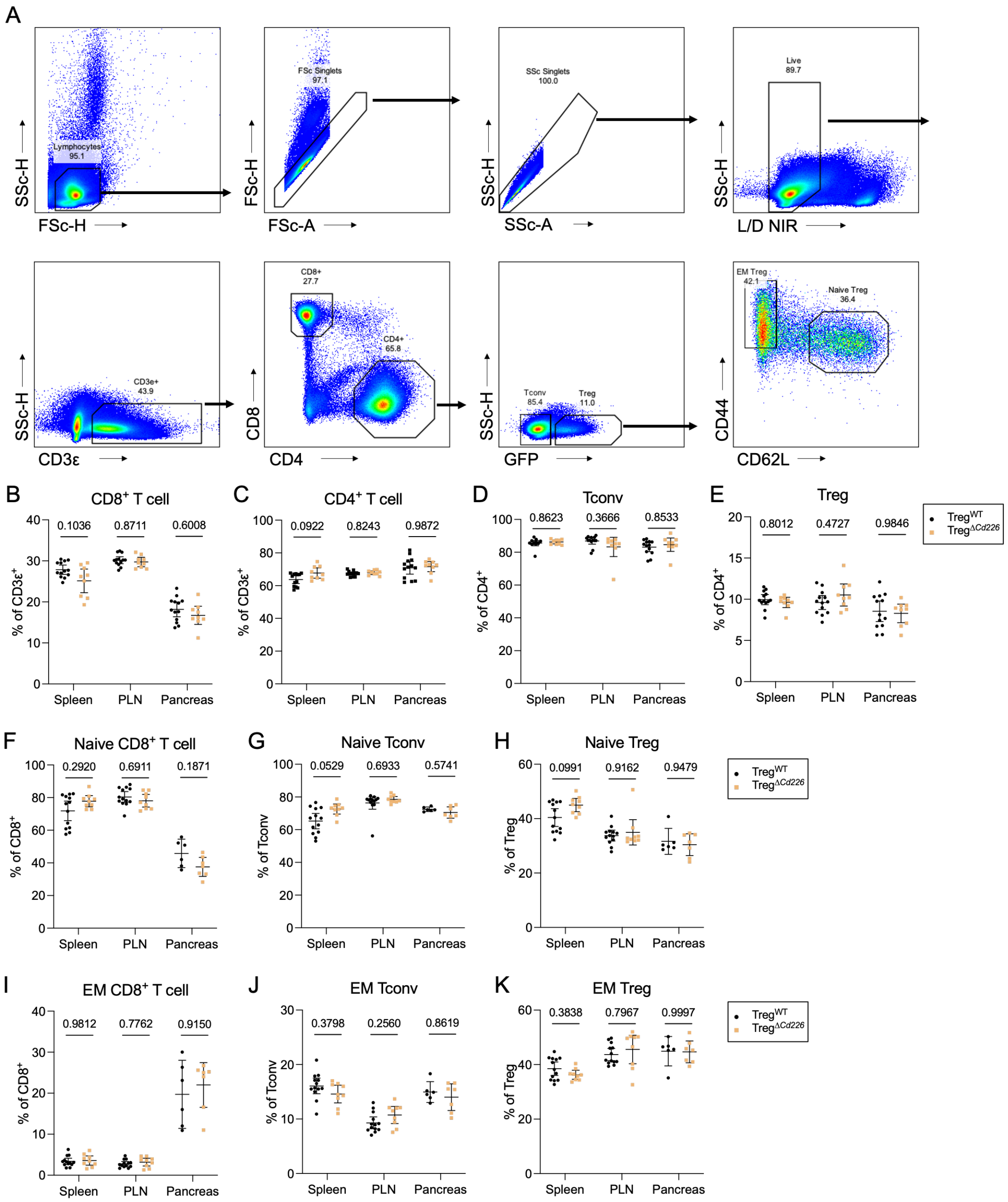


**Supplemental Figure 3. CD8$^+$ T cell, Tconv, and Treg subset frequencies are maintained in Treg$^{\Delta Cd226}$ mice. (A)** Example flow cytometry gating scheme for the cKO strain. Quantification of percentages of **(B)** CD8$^+$ and **(C)** CD4$^+$ T cells as well as **(D)** CD4$^+$ Tconv and **(E)** Tregs in the spleens, PLNs, and pancreata of 12-week-old female mice. **(F-H)** Quantification of CD44$^-$CD62L$^+$ naïve CD8$^+$, CD4$^+$ Tconv, and Tregs in the spleen, PLN, and pancreas of 12-week-old female

Treg$^{WT}$ (*n*=13) and Treg$^{\Delta Cd226}$ (*n*=9) mice. **(I-K)** Quantification of CD44$^{+}$CD62L$^{-}$ EM CD8$^{+}$, CD4$^{+}$ Tconv, and Tregs in the spleen, PLN, and pancreas. Statistical analyses were performed with multiple t-tests with Sidak-Bonferroni multiple comparisons correction.

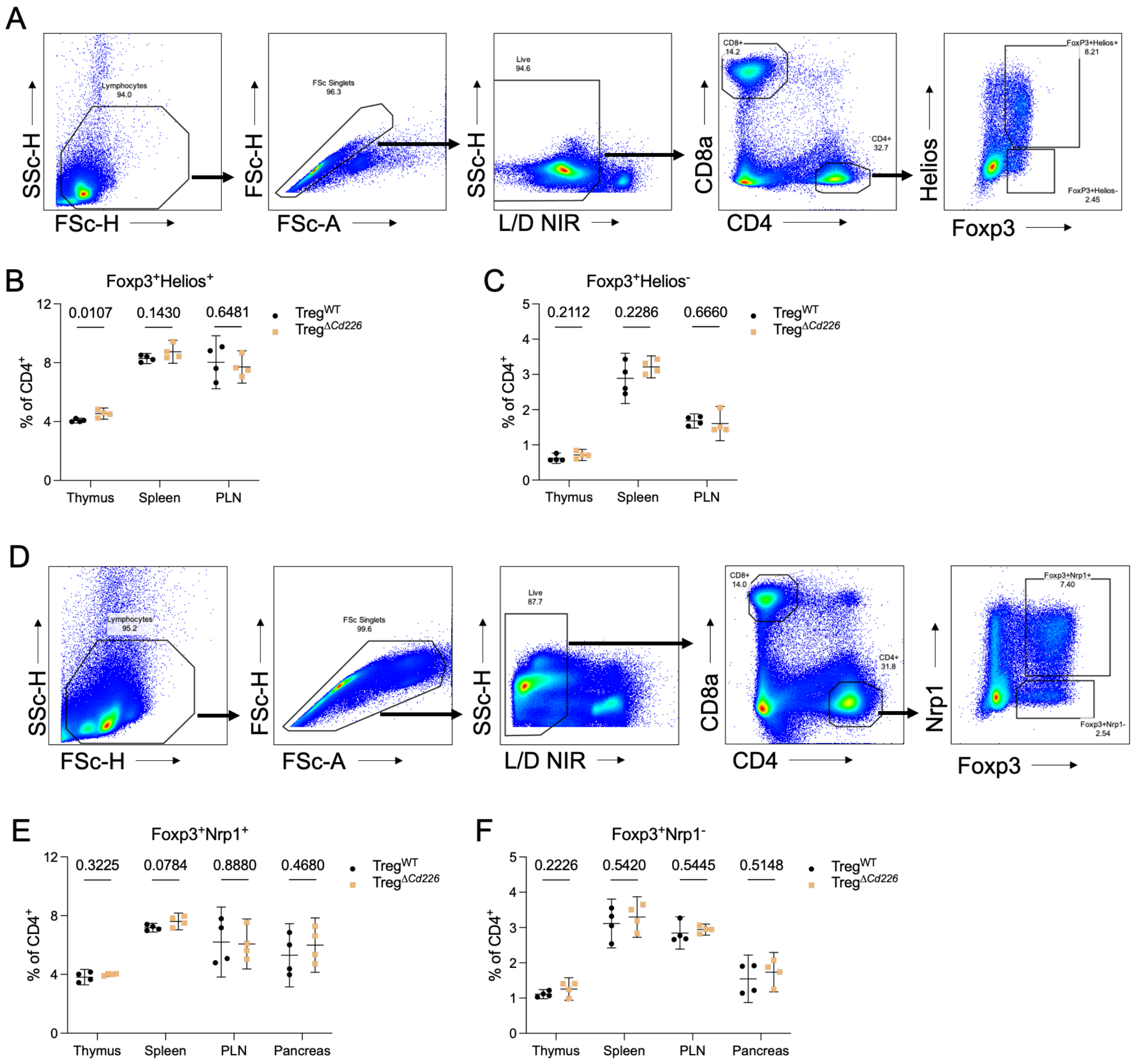


**Supplemental Figure 4. Natural or adaptive Treg frequencies are unaffected by *Cd226* cKO.** **(A)** Standard flow cytometric gating scheme for Tregs with representative data shown from spleen. Percentages of parent population are shown underneath the gate names. **(B-C)** Quantification of Foxp3$^+$Helios$^+$ and Foxp3$^+$Helios$^-$ Tregs in the thymus, spleen, and PLN of 12-week-old pre-diabetic female Treg$^{WT}$ (black) and Treg$^{\Delta Cd226}$ (orange) mice. **(D)** Standard flow cytometric gating scheme with representative data shown from spleen **(E-F)** Quantification of Foxp3$^+$Nrp1$^+$ and Foxp3$^+$Nrp1$^-$ Tregs in the thymus, spleen, PLN, and pancreas of Treg$^{WT}$ (black) and Treg$^{\Delta Cd226}$ (orange) mice. P values shown on figure, two-way ANOVA ($n$=4 biological replicates).
